# Elucidating the patterns of pleiotropy and its biological relevance in maize

**DOI:** 10.1101/2022.07.20.500810

**Authors:** Merritt Khaipho-Burch, Taylor Ferebee, Anju Giri, Guillaume Ramstein, Brandon Monier, Emily Yi, M. Cinta Romay, Edward S. Buckler

**Author notes:** Corresponding author: Merritt Khaipho-Burch.

## Abstract

Pleiotropy - when a single gene controls two or more seemingly unrelated traits - has been shown to impact genes with effects on flowering time, leaf architecture, and inflorescence morphology in maize. However, the genome-wide impact of true biological pleiotropy across all maize phenotypes is largely unknown. Here we investigate the extent to which biological pleiotropy impacts phenotypes within maize through GWAS summary statistics reanalyzed from previously published metabolite, field, and expression phenotypes across the Nested Association Mapping population and Goodman Association Panel. Through phenotypic saturation of 120,597 traits, we obtain over 480 million significant quantitative trait nucleotides. We estimate that only 1.56-32.3% of intervals show some degree of pleiotropy. We then assessed the relationship between pleiotropy and various biological features such as gene expression, chromatin accessibility, sequence conservation, and enrichment for gene ontology terms. We find very little relationship between pleiotropy and these variables when compared to permuted pleiotropy. We hypothesize that biological pleiotropy of common alleles is not widespread in maize and is highly impacted by nuisance terms such as population structure and linkage disequilibrium. Natural selection on large standing natural variation in maize populations may target wide- and large-effect variants, leaving the prevalence of detectable pleiotropy relatively low.

**Author Summary:** The genetic basis of complex traits has been thought to exhibit pleiotropy, which is the notion that a single locus can control two or more unrelated traits. Widespread reports in the human disease literature show genomic signatures of pleiotropic loci across many traits. However, little is known about the prevalence and behavior of pleiotropy in maize across a large number of phenotypes. Using association mapping of common alleles in over one hundred thousand traits, we determine how pleiotropic each region was and use these quantitative scores to functionally characterize each region of the genome. Our results show little evidence that pleiotropy is a common phenomenon in maize. We observed that maize does not exhibit the same pleiotropic characteristics as human diseases in terms of prevalence, gene expression, chromatin accessibility, or sequence conservation. Rather than pervasive pleiotropy, we hypothesize that strong selection on large and wide effect loci and the need for trait independence at the gene level keep the prevalence of pleiotropy low, thus, allowing for the adaptation of maize varieties to novel environments and conditions.

## Introduction

Pleiotropy, a term defined by German geneticist Ludwig Plate in 1910, refers to the phenomenon where a single locus can affect two or more seemingly unrelated traits [1]. Biological pleiotropy (also known as horizontal pleiotropy) describes situations in which a single causal variant affects two or more traits, different causal variants colocalize in the same gene and are tagged by the same genetic variant, or different causal variants colocalize in the same gene and are tagged by different genetic variants [2]. In contrast, mediated pleiotropy (also known as vertical pleiotropy) occurs when one variant impacts a phenotype that is then causal for another phenotype. Instances of mediated pleiotropy are difficult to ascertain and require a deeper understanding of how different biological pathways interact. Both horizontal and vertical pleiotropy can be biased by spurious pleiotropy arising from misclassifying relationships between traits or high linkage disequilibrium tagging nearby, non-causal variants [2].

Some examples of single-locus pleiotropy have been observed in maize. The *Vegetative to generative transition 1* (*Vgt1*) regulatory sequence controls leaf number and flowering time by regulating an *APETALA2*-like gene, *ZmRap2.7*, through what may be mediated pleiotropy [3–6]. Polymorphisms within *Vgt1* cause an additive genetic effect of roughly 2-4.5 days to pollen shed and an increase in the number of internodes. True pleiotropy of this locus was confirmed through introgression of the *Vgt1* locus into transgenic plants [7,8]. The *teosinte branched 1* (*tb1*) locus is an example of biological pleiotropy, which impacts the number and length of internodes in lateral branches and inflorescences due to the upregulation of gene expression in maize compared to teosinte. This upregulation results from the insertion of a *Hopscotch* transposable element 64 kb upstream of *tb1 [9–12]. Fine-mapping of tb1* confirmed this biological pleiotropy [10,13,14]. Although other examples of biological pleiotropy at single genes have been described in maize, many have not been confirmed through fine-mapping to distinguish between tight linkage and true pleiotropy [15].

With joint linkage studies and genome-wide association studies (GWAS) in plants, the search for pleiotropy became more promising. However, few maize genes have shown true biological or mediated pleiotropy. Some pleiotropy is present between phenotypes of the same trait type, like the architecture of tassels, ears, leaves, or flowering time; however, very few single nucleotide polymorphisms (SNPs) are shared between different trait types [16,17]. Some evidence of pleiotropy between root, leaf, and flowering traits has been shown but requires additional testing and validation [17]. A similar maize study showed evidence of pleiotropy but within inflorescence traits, such as kernel row number with thousand kernel weight and tassel standing, tassel length, and tassel spike length [18]. The only robust examples of pleiotropy discovered in maize are between flowering traits and leaf number [16,18]. The overlap of only a few key genes between flowering and leaf number agree with prior literature in that both traits share a developmental pathway and that the transition from halting leaf growth to the initiation of tassel development is highly correlated [3,19–21].

A lack of pervasive pleiotropy was also observed in sorghum, where a large set of 234 plant architecture and agronomic traits was investigated within the Sorghum Association Panel [22]. The lack of marker saturation within GWAS models may have contributed to these results, or potentially relying on Bayesian SNP selection models to consistently select the same pleiotropic SNP across multiple independent traits. However, even with these methods, some large-effect pleiotropic loci were found along with novel pleiotropic sites between traits [22]. The lack of additional shared genes between different trait types within prior studies suggests that pleiotropy is not a common phenomenon in maize or sorghum. Many traits showing this apparent pleiotropy may be under similar regulation due to the traits sharing the same developmental phytomers. These developmentally correlated traits which share high morphogenic relatedness may signal the illusion that pleiotropy is much more widespread than it is.

Some studies have calculated principal components over their traits to create uncorrelated variables used to determine the degree of pleiotropy [18]. Using uncorrelated traits addresses the concern that all phenotypes are inherently related and not truly distinct if they share the same evolutionary or developmental origin [23]. However, many of these studies may miss large, causal loci impacting their phenotypes of interest in principal-component based traits as opposed to observed trait values, biasing estimates of pleiotropy [18]. Additionally, GWAS-based pleiotropy studies have been restricted to finding common alleles with modest to large effects, while rare variants with large effects and common variants with small effects are difficult to find [24–27]. With a small range of allelic detection, the ability to discern all putatively pleiotropic effects is limited. This may be partially responsible for the minimal pleiotropy described in crop plants [18,22].

Pleiotropy has been described much more frequently in human disease literature. Some studies reported that nearly half of the disease-associated genes within the human GWAS Catalogue were pleiotropic [27]. Similar pleiotropy was found among other human traits and disease studies [28–30]. To characterize pleiotropic loci, a study investigating 321 vertebrate genes showed that highly pleiotropic genes showed higher gene expression [31]. In human studies, pleiotropic genes were strongly associated with higher expression [28], were expressed in more tissues [29], and were enriched in active chromatin states compared to genome-wide loci [29]. Additionally, major pleiotropic loci in vertebrate genes evolve at slower rates than less pleiotropic genes, suggesting that the degree of pleiotropy may constrain adaptive evolution [31].

While human studies are more likely to be looking for pleiotropy within diseases, the extent of pleiotropy among common adaptive variants in maize remains elusive. Although instances of pleiotropy have been described in maize, many of these studies have been limited to large-effect domestication loci [7,10,14], in the number of traits they investigate [18,32,33], low mapping resolution of QTL [3,16,34,35], or have variable methods of distinguishing and selecting a true biologically pleiotropic locus [36]. Additionally, a major limitation to pleiotropy studies in maize is investigator bias in phenotyping, relatively common, easier to measure field-based traits that may behave differently from molecular phenotypes such as gene expression or metabolites. These field versus molecular phenotypes may show distinct patterns of pleiotropy that have been relatively unexplored in the literature. Although developmentally, some of these bulked traits may share the same generating phytomers, they still offer an alternative view of pleiotropy.

In this study, we investigated the pervasiveness of pleiotropy in maize using a large set of 120,597 previously published phenotypes in the US Nested Association Mapping population and the Goodman Association Panel (hereafter NAM and GAP, respectively) that generated 480 million significant quantitative trait nucleotides. We used these association results to calculate a quantitative estimate of how pleiotropic each genic and intergenic region was within the genome. Then, we functionally characterized these loci through gene expression, chromatin openness, and sequence conservation. We aimed to investigate instances of true biological pleiotropy where variants directly impact multiple traits while trying to limit the amount of spurious pleiotropy occurring from linkage disequilibrium.

## Results

### Genome-wide association of traits across NAM and GAP

To determine the extent of pleiotropy within the maize genome, we performed GWAS on 120,597 traits across NAM (n = 5186) and GAP (n = 271). These traits are outlined in further detail in Supplemental Tables 1-3. These traits included 121 NAM field traits, 222 GAP field traits (see methods), 3,873 GAP leaf metabolite mass features [37], and 116,381 GAP expression values across seven diverse tissues [38]. The traits were categorized into four categories based on the population and type of trait measured. These population-trait categories were: 1) NAM field, 2) GAP field, 3) GAP mass features, and 4) GAP expression traits (S1 Table). A schematic of our methods is shown in Fig 1. Genotype data came from HapMap 3.2.1 SNPs and were imputed using Beagle 5.1 with the USDA-ARS NCRPIS collection SNPs as the reference. After filtering, roughly 15 million NAM SNPs and 17 million GAP SNPs remained for association mapping.

**Fig 1:**
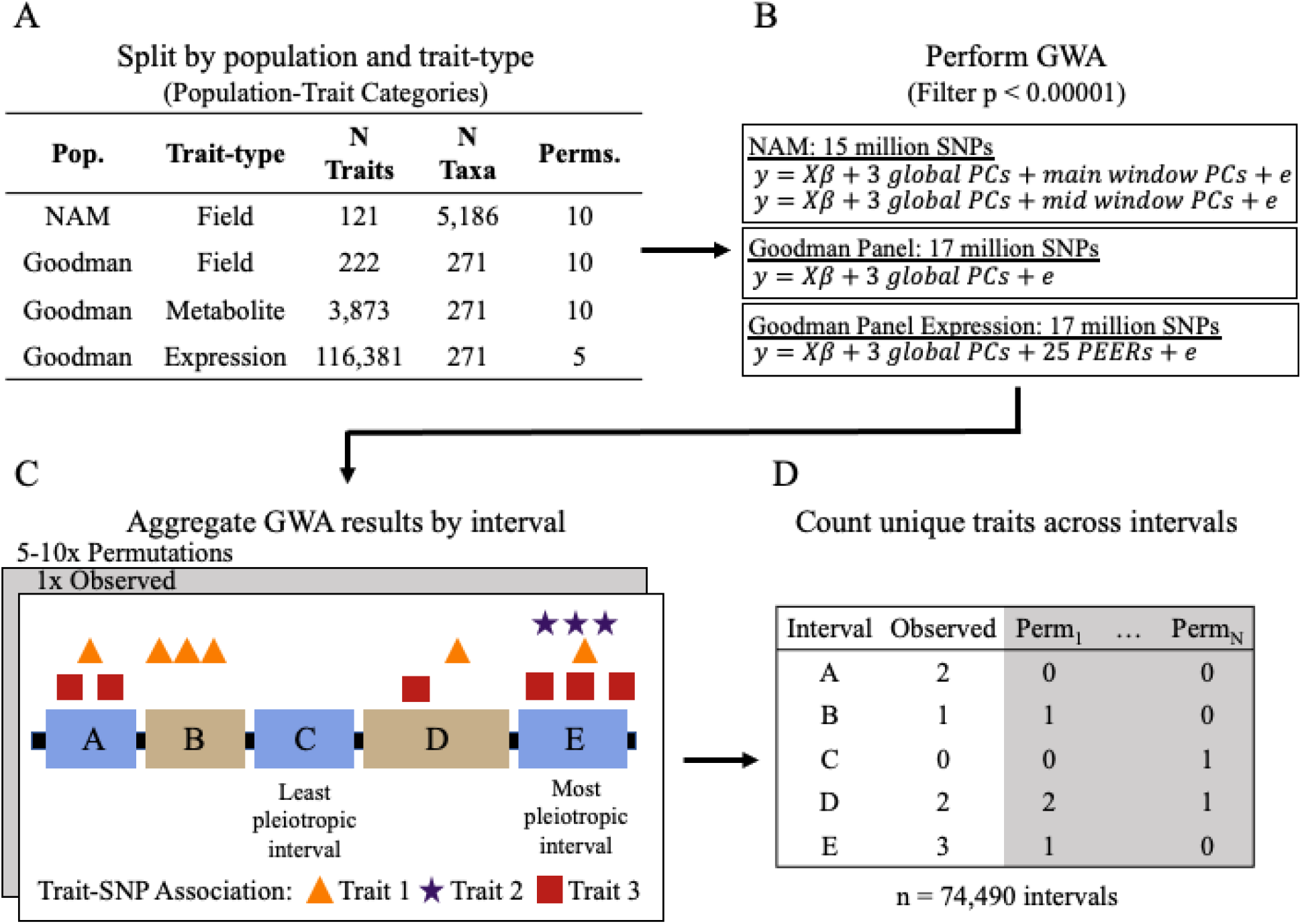
Schematic of the methods used within this study. (A) A total of 120,597 traits were gathered across four population-trait categories: 1) NAM field, 2) GAP field, 3) GAP mass features, and 4) GAP expression. Trait values were shuffled randomly (i.e., permuted) five or ten times to create null distributions of no pleiotropy. (B) The observed and permuted data were then mapped using the Fast Association method in rTASSEL using 15 million NAM SNPs and 17 million GAP SNPs using a combination of global and window principal components and PEER factors to control population structure and experimental design. All results with p < 0.00001 were kept for both the permuted and observed data. (C) GWAS results were then split by trait type and permutation and were then aggregated into genic and intergenic intervals based on the the significant B73 v4 trait-SNP association coordinate. (D) A matrix of the number of unique traits mapping to each interval was used to estimate biological and mediated pleiotropy and was collected independently for each population-trait type.

### The prevalence of pleiotropy is low but shows variation by trait type and population

Each population-trait type was investigated independently due to the uniqueness of the genetic architecture of each population, the variable trait sample size, and to investigate the differences in the patterns of pleiotropy between trait types. For each population-trait type, the number of unique traits mapping to each genic and intergenic interval was counted for both the observed and permuted association results to gain a quantitative metric on how pleiotropic each interval was within the genome. These pleiotropy estimates include biological, mediated, and spurious pleiotropy, as distinguishing between the different forms is difficult. A total of 75,490 intervals with a mean length of 27,902 bp were used. Counts for the null distributions of no pleiotropy originated from ten permutations of the field and mass features, while the expression phenotypes were permuted five times. We adjusted all of our permuted phenotypes before GWA by the same principal components or PEER factors used within the GWA models to not inflate p-values due to the underlying population structure held within principal components.

We compared our distribution of observed significant traits per interval to the permuted trait counts. Across populations and trait types, 67.7-98.4% of intervals have 0-1 significant traits associated with them (Fig 2). Most intervals in the observed data controlled more traits than the permutations. This does not completely imply pervasive pleiotropy but rather, most traits are polygenic.

**Fig 2:**
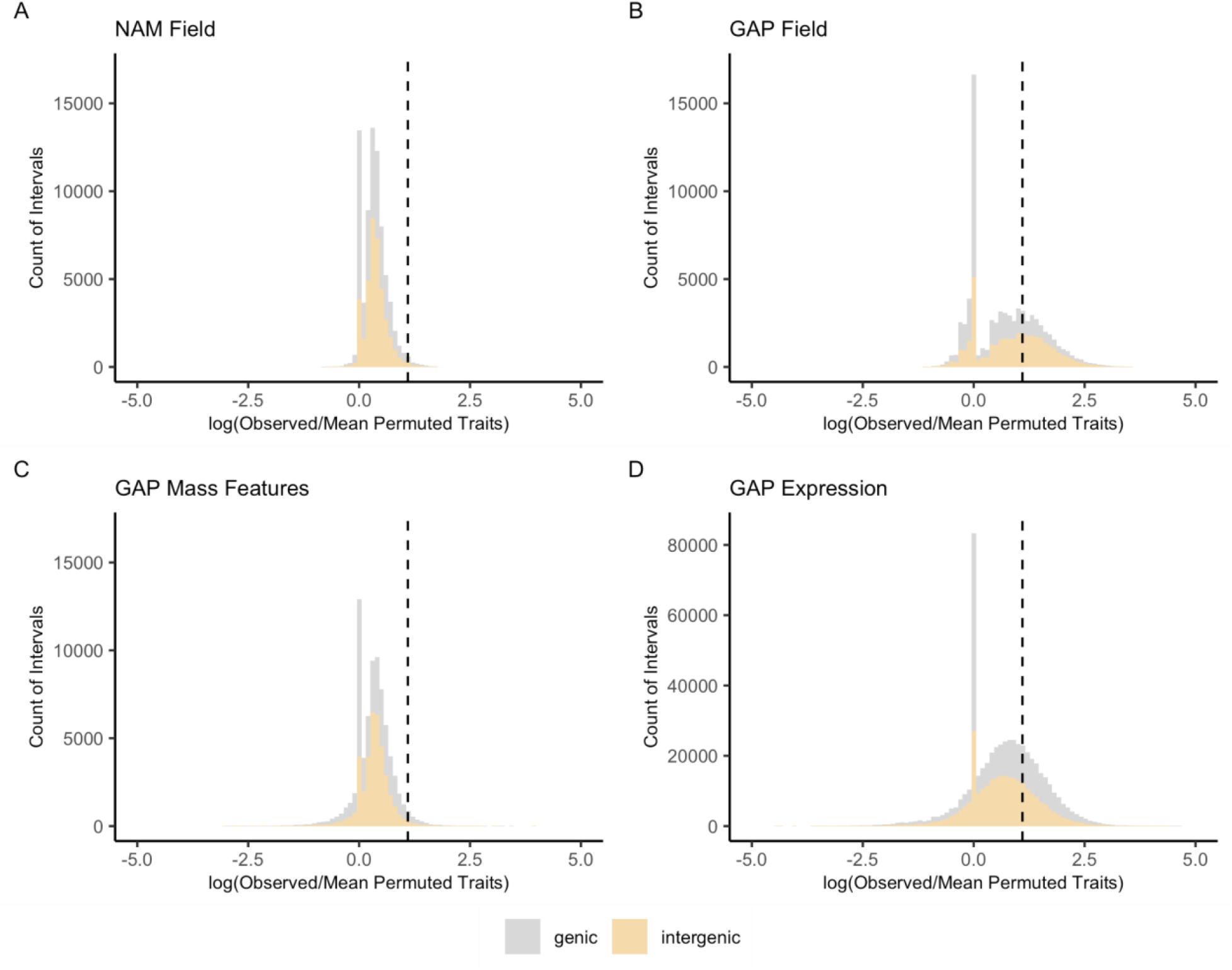
Distribution of QTL by interval. Of the intervals showing a five-fold higher proportion of pleiotropy over their permutations (right of the vertical dashed line), intergenic ranges tend to exhibit higher pleiotropy than genic ranges. Each value along the x-axis was calculated from the natural log of the number of observed traits mapping to each interval divided by the mean count of traits in the permuted data with a pseudo-count of plus one in the numerator and denominator. Values right of the vertical dashed line indicate higher pleiotropy in the permuted data versus the observed data suggesting the prevalence of high noise or no trait-SNP associations in either the observed or permuted data (peak at zero). Distributions are split into genic (gray) and intergenic (yellow) intervals for (a) NAM field, (b) GAP field, (c) GAP mass features, and (d) GAP expression traits.

For intervals showing five-fold higher evidence in the observed versus the mean permuted data, NAM field phenotypes show some pleiotropy at intervals putatively associated with 2-19 traits (median = 5 traits) in 1.56% of intervals. GAP field traits show pleiotropy associated with 2-65 traits (median = 9) in 32.3% of intervals genome-wide. GAP mass features showed much less pleiotropy, with 3.03% of intervals associating with between 2-695 traits (median = 63). Lastly, GAP expression data showed slightly more pleiotropy with 31% of intervals associating with 2-6264 traits (median = 391). In relation to the total number of expression traits (n = 116,381), a median of 0.33% of all expression traits impacted these highly pleiotropic intervals. For GAP mass features (n = 3,873), a median of 1.62% of all mass features contributed to these high pleiotropy estimates. Overall this suggests that there may be a set of genes with high pleiotropy, but as we will show below, this is mostly a product of factors impacting mapping.

These pleiotropy estimates for GAP field and expression data may be higher than NAM field and GAP mass feature estimates. For GAP field data, these pleiotropy estimates might be slightly inflated due to residual population structure that was not controlled for well during the association mapping step [39], but mapped this way for consistency with metabolite and expression data). Additionally, for all the population and trait types, retaining all trait-SNP associations with p < 0.00001 may have introduced noise depending on the genetic architecture of the trait being mapped that could, in turn, overrepresent the amount of pleiotropy present.

### Mapping features explain most pleiotropy

We analyzed various biological and nuisance (GWAS mapping) features to understand their importance in explaining this apparent pleiotropy. We evaluated two hypotheses that this apparent pleiotropy was due to characteristics of genetic mapping or various aspects of biological regulation and mutation-selection balance. Biological features consisted of the maximum gene and protein expression values in B73 across twenty-three diverse tissues [40], the mean Genomic Evolutionary Rate Profiling (GERP) scores at sites overlapping with significant GWA hits [41], and the number of ATAC-seq peaks [42]. Nuisance features, which have to do with genetic mapping and resolution, were similarly modeled and included the average R^2^ linkage disequilibrium within NAM and GAP, the total number of SNPs per interval used for the GWA analysis, and interval size. We also included an adjusted GAP pleiotropy estimate for NAM field, GAP field, and GAP mass feature models to see if pleiotropy at an intermediate molecular step had any effect on field or metabolite regulation.

All pleiotropy scores are highly correlated with nuisance terms (Fig 3). The number of input SNPs, the interval size, and average linkage disequilibrium all show strong, positive correlations with pleiotropy values. Additionally, pleiotropy values between populations and trait types are highly correlated with each other. For the biological features, there is a weak correlation between pleiotropy and GERP, max protein expression, max RNA expression, and the number of ATACseq peaks. A full correlation matrix including gene ontology (GO) terms is available in S1 Fig.

**Fig 3:**
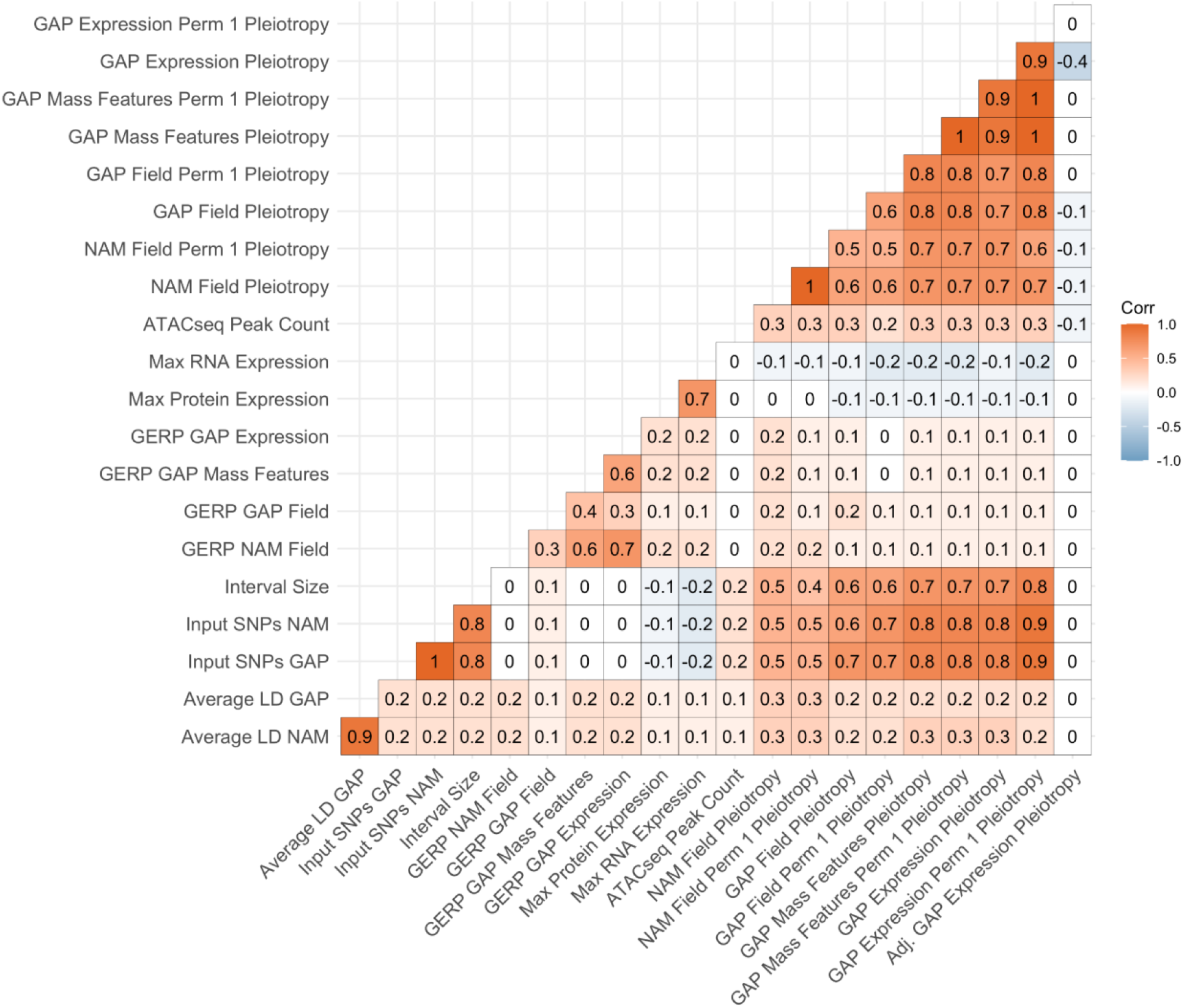
Correlation matrix of key biological, nuisance (mapping), and pleiotropy values. Values indicate the Pearson correlation between values colored by the strength of their correlation.

With these biological, nuisance, and adjusted pleiotropy terms we ran four models (Models 1-4) that varied in their input features and model architectures. Each model type was run eight separate times for each population-trait type (NAM field, GAP field, GAP mass features, and GAP expression) and permutation type (observed versus permuted data). All models were trained using chromosomes 1–9 and tested using chromosome 10 interval data. Model 1 was a random forest model that included only nuisance, or mapping terms. Model accuracy varied by the population and trait type (R^2^ = 0.41-0.84, S5 Table). Prediction on held-out chromosome 10 data showed a strong relationship between predicted and observed pleiotropy (R^2^ = 0.92-0.98, Fig 4), suggesting that pleiotropy within these populations can be predominantly described by mapping terms.

**Fig 4:**
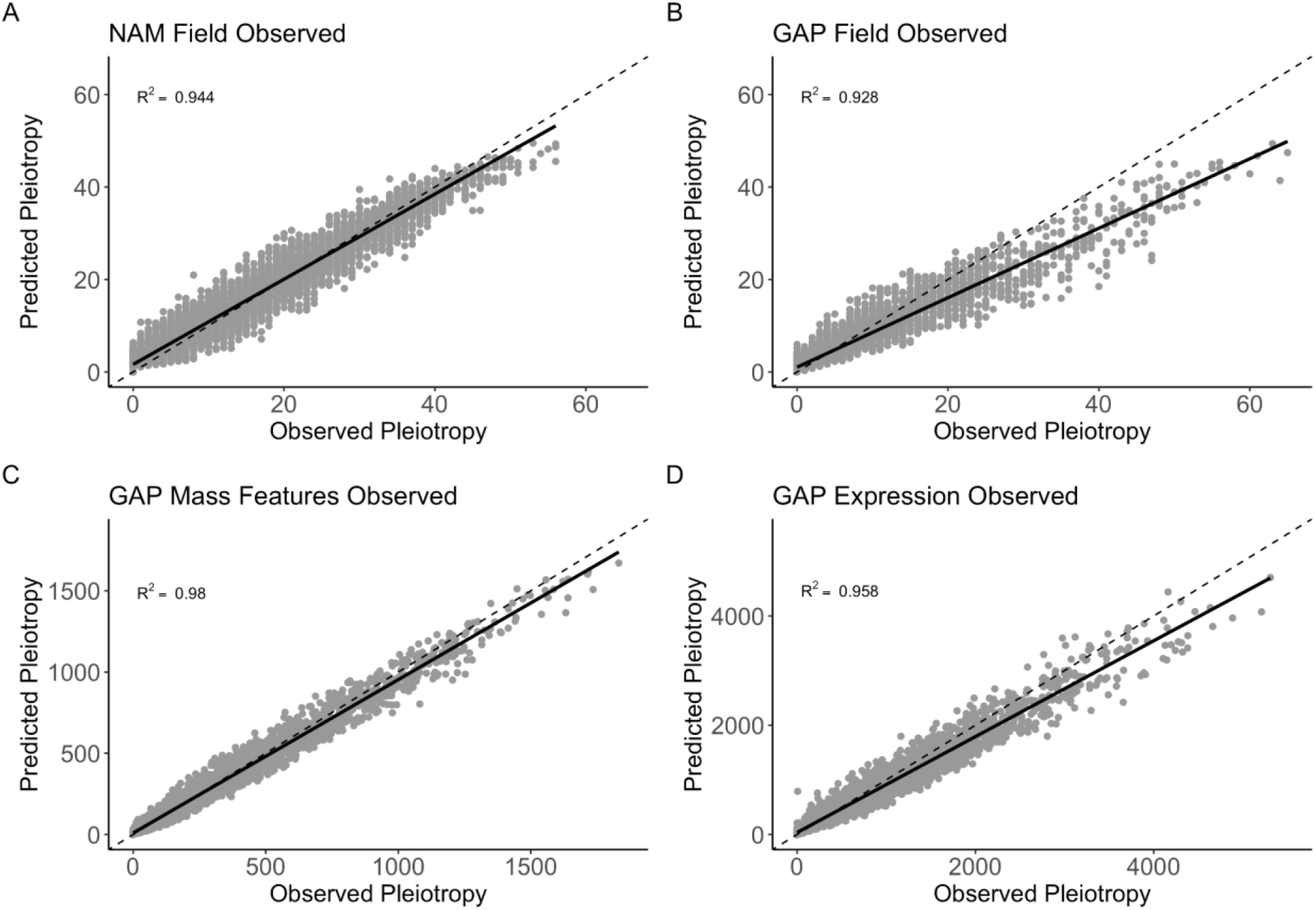
Predicted versus observed pleiotropy values for Model 1 (random forest with only nuisance terms) from four individual random forest models trained on observed pleiotropy values from chromosomes 1-9 and tested on chromosome 10 data. The dashed line represents the 1-1 identity line, while the solid line represents fitted values. Panels show the results for the (a) NAM field, (b) GAP field, (c) GAP mass features, and (d) GAP expression data.

Model 2 was trained on pleiotropy scores using all biological and nuisance terms described previously using a random forest model. Overall, the models performed well, with R^2^ values ranging from 0.439 to 0.871 (S5 Table), and showed good performance when predicting chromosome 10 data across models (S3 Fig). Across population-trait types in the observed and permuted data, the most important terms in predicting pleiotropy were nuisance variables that included the number of input SNPs, average R^2^ linkage disequilibrium, and interval size (Fig 3). All biological features had lower importance for predicting pleiotropy than the nuisance variables. The mean GERP score showed moderate importance across models, and RNA and protein expression showed the least importance of these biological features. All other biological features were only moderately important. The ranking of features between the observed and permuted data models remained largely the same with very little deviation (S2 Fig).

Model 3 was also a random forest model trained using the same biological and nuisance terms as Model 2 but included fourteen gene ontology terms thought to be important in explaining highly pleiotropic loci. Model 4 was a gradient boosting model with only the terms described in Model 2 (no gene ontology terms). Models 2, 3, and 4 performed similarly, with the nuisance variables showing the highest relative importance (Fig 5, S2, and S6 Figs). All models had comparable model performance, with the gradient boosting method (Model 4) having slightly higher prediction accuracy (Model 4 R^2^ 0.627-0.913) over the random forest models (Model 3 R^2^ 0.332-0.987; S3, S5, S7 Figs, and S5 Table). For Model 3, which included gene ontology terms, all gene ontology terms showed lower relative importance than the biological and nuisance variables (S4 Fig).

**Fig 5:**
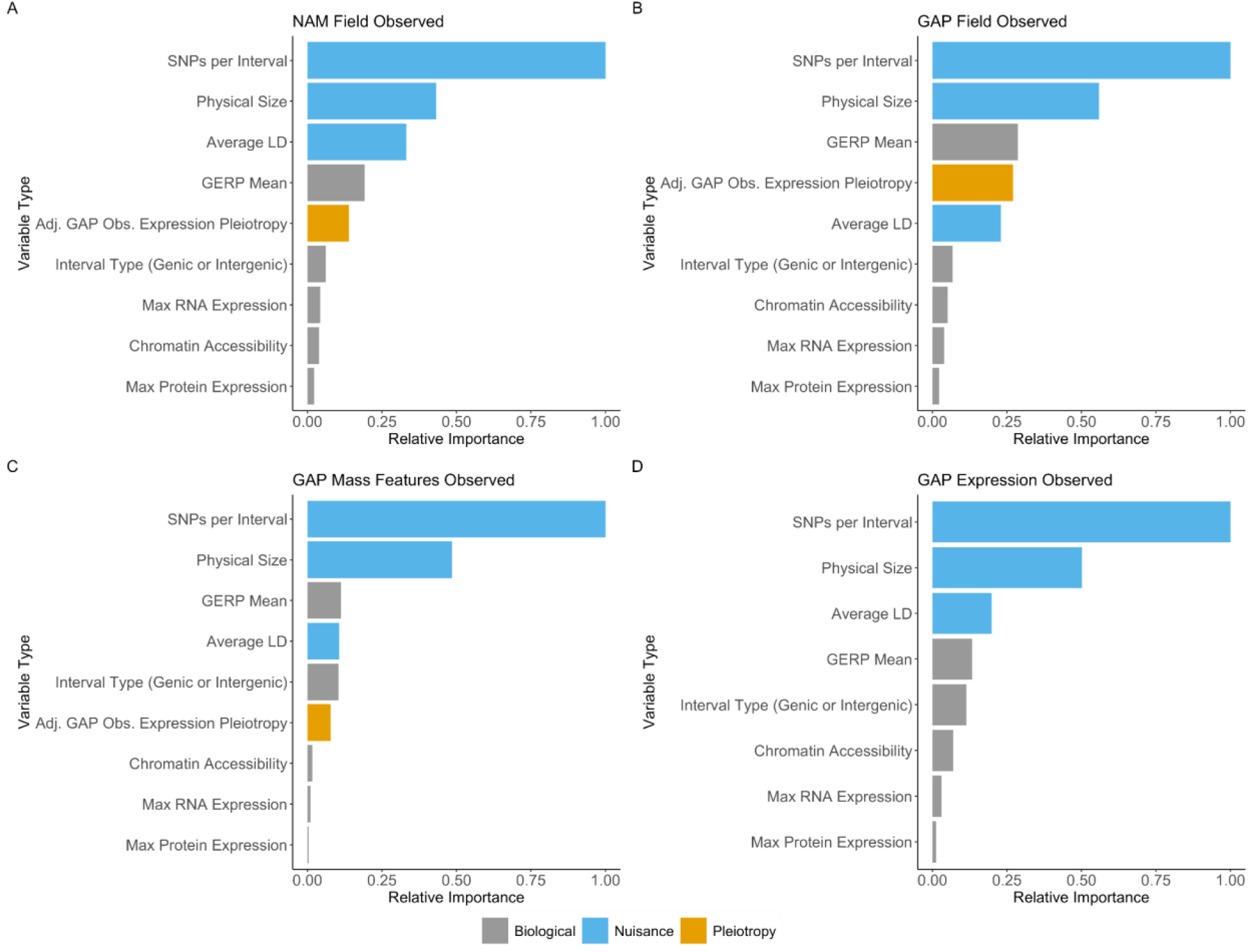
Relative importance scores from random forest models suggest high importance for nuisance terms and low importance for biologically relevant terms in explaining pleiotropy. The random forest results shown are from Model 2 (with no gene-ontology terms). Across all four population-trait categories, nuisance variables such as the number of input SNPs, interval size, and average linkage-disequilibrium showed higher relative importance over biological features. The plots show data for the observed (not permuted) data for (a) NAM field data, (b) GAP field data, (c) GAP mass features, and (d) GAP expression data. Model 2 results for the permuted data are available in S2 Fig.

Due to the overwhelmingly large effect of mapping features in describing pleiotropy, an adjusted pleiotropy statistic was created. This adjusted statistic was created from Model 1 by subtracting the predicted pleiotropy from the observed pleiotropy score and was used to look at different biological relationships in more depth. The relationship between adjusted pleiotropy and observed pleiotropy is available in S8 Fig. We then regressed these adjusted pleiotropy values against the max RNA expression across 23 diverse B73 tissues and the mean GERP score. We found no difference in the relationship between adjusted pleiotropy with GERP or the max RNA expression value for NAM field, GAP field, or GAP mass features compared to the permuted data. However, we did find a change in the relationship between RNA expression and GERP with GAP expression pleiotropy data in the observed versus the permuted data, where more pleiotropic intervals showed higher GERP sequence conservation and lower expression values (S9 Fig, S10 Fig). These relationships of observed GAP expression pleiotropy with max RNA expression (adj. R^2^ = -2.41e-05, p = 0.767) and GERP (adj. R^2^ = -6.68e-06, p = 0.4016) were not significant.

With these adjusted pleiotropy scores, we then investigated the most pleiotropic genes within each population and trait type. The top twenty most pleiotropic genes by population and trait type are presented in S7 Table. Genes that were shared across populations and traits types included *Zm00001d035776*, which is a paired amphipathic helix protein shown to be involved in regulating flowering time and internode number in Arabidopsis [43], pleiotropic in both GAP field and expression data. *Zm00001d040920* is a tRNA methyltransferase pleiotropic within NAM field and GAP expression data. Lastly, we searched for overlap in our top twenty highly pleiotropic genes within a prior large-scale maize pleiotropy study [17]. We found one shared sphingomyelin synthetase family protein (*Zm00001d034600*), in GAP field traits that impacted two tassel traits, and a G2-like1 gene (*Zm00001d044785*) in GAP mass features that impacted six root traits <u>(Mural et al. 2022)</u>.

### Gene ontology suggests differential enrichment for highly and lowly pleiotropic intervals

To determine if there was any differential enrichment in gene ontology terms among our putatively pleiotropic sites, we categorized our genic intervals into highly and lowly pleiotropic sets based on their adjusted pleiotropy values. Genic intervals were classified as lowly or highly pleiotropic if they had the lowest or highest twenty-five percent of above zero adjusted pleiotropy values within each population-trait type. We then queried how enriched the lowly and highly pleiotropic gene sets were for molecular function, biological process, and cellular component terms. No cellular component terms were significant.

For molecular function, we found an enrichment of highly pleiotropic terms for small molecule binding, and ribonucleotide binding for GAP field data, GAP mass features, and GAP expression data (Fig 6). Additionally, terms involving tubulin binding, transferase activity, potassium ion binding, and ligase activity were shown to be significant in highly pleiotropic intervals in some populations. For intervals with low pleiotropy, terms associated with transcription regulator activity were significant for GAP field, GAP mass features, and GAP expression data.

**Fig 6:**
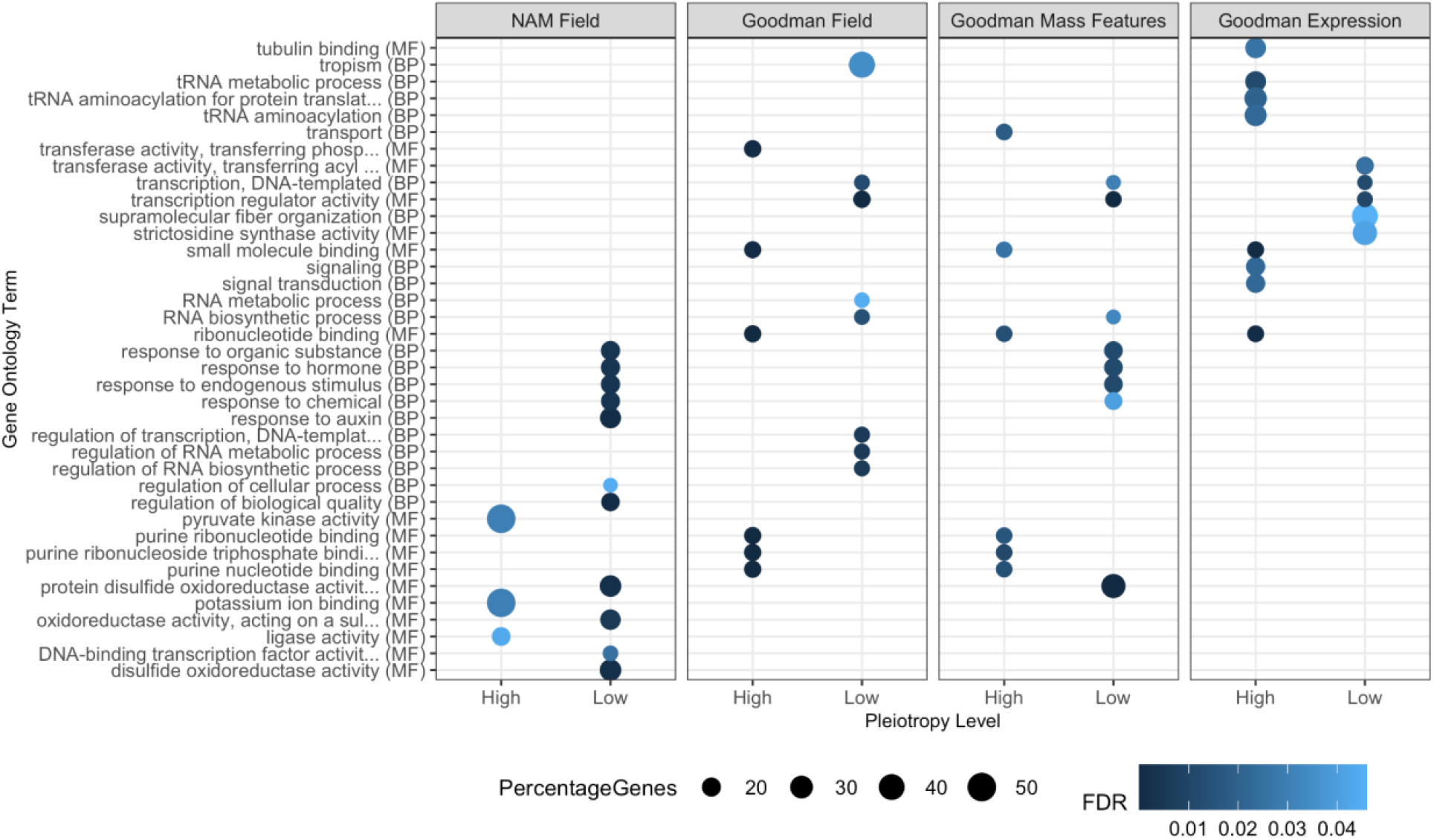
Gene ontology enrichment for highly and lowly pleiotropic intervals for the four population-trait categories. Highly and lowly pleiotropic values fell within the top and bottom 25th percentile of adjusted pleiotropy values for each population-trait type. All molecular function, biological process, and cellular component terms were tested. Terms with false discovery rate (FDR) corrected p-values below p = 0.05 were retained. The x-axis represents highly or lowly pleiotropic intervals split by the population and trait type, and the y-axis has the top significant gene ontology terms. The value of the FDR significance level is colored in blue and the size of the dots represents the proportion of genes found in that GO category.

For biological process terms, we found an enrichment of lowly pleiotropic terms for tropism, RNA metabolic processes, response to auxin, regulation of DNA transcription, and regulation of RNA metabolic and biosynthetic processes were shown to be significant in some populations. Lowly pleiotropic terms that were significant between populations were responses to organic substances, hormones, stimulus, and chemicals. Additional lowly pleiotropic terms shared between populations were DNA templated transcription, and RNA biosynthetic processes. Highly pleiotropic biological processes included terms involved with transport in GAP mass features.

## Discussion

Our results showed modest but pervasive pleiotropy in the Nested Association Mapping and Goodman Association Panel across a wide range of field, metabolite, and expression phenotypes that could almost always be highly explained by mapping resolution characteristics. Across 120,597 maize traits, we showed that some apparent pleiotropy is present across populations, however, much of this pleiotropy is highly correlated and explained by association mapping parameters in multiple random forest models. A total of 67.7-98.4% of intervals were associated with zero to one traits. These distributions are reminiscent of the L-shaped pleiotropy distributions derived from GWA- and QTL-based studies in other species like mouse, yeast, nematode, and stickleback fish, where the large majority of loci have little to no effect on traits while a minimal subset of sites show pleiotropic effects [44,45]. Within these distributions, intergenic intervals had higher pleiotropy values confirming prior results in maize where most GWA hits fall within larger, intergenic regions rather than genic regions [46], which would lead to higher overall pleiotropy scores. One caveat to this study is that we cannot detect pleiotropy in fixed mutations because our estimates are based on the prevalence and frequency of association results–values dependent on allele frequencies within populations. We are thus more likely to find common variants with only modest effects on phenotypes than rare variants with small effects [24–26].

Given our distribution of pleiotropic loci derived from common alleles, we investigated the top twenty intervals with the largest adjusted pleiotropy values. Found across two populations was a paired amphipathic helix protein (*Zm00001d035776*) shown to be involved in regulating flowering time and internode number in Arabidopsis [43] and a tRNA methyltransferase (*Zm00001d040920*). All other genes within, or next to the most pleiotropic intervals, were only present in one population. In GAP field traits, *xylanase/glycosyl hydrolase10* (*xyn10, Zm00001d039958*) has been shown to impact mesocotyl length after deep-seeding [47], with the larger family of glycoside hydrolases being involved in plant cell enlargement and composition [48]. A highly pleiotropic locus in GAP mass features included *empty pericarp2* (*Zm00001d005675*), which is involved in embryo morphogenesis, shoot development, and the negative regulation of the heat shock response [49–51]. GAP expression data showed *ZIM-transcription factor 8* (*zim8, Zm00001d004277*), a candidate gene for *Striga* damage [52], and kernel composition and popping expansion [53].

To understand the genetic architecture behind these pleiotropic loci we ran multiple random forest and gradient tree boosting models investigating the relationship between biological and nuisance (mapping) variables to characterize the patterns of pleiotropy. Nuisance variables, including the interval size, average linkage disequilibrium, and the number of input SNPs, showed the highest relative importance across models. Of much lower relative importance were biological features, including maximum gene expression across multiple B73 tissues [40], the amount of accessible chromatin [42], and the average evolutionary conservation [41]. Even after adjusting pleiotropy scores using the nuisance variables, we found no strong or significant relationship with maximum gene expression or the average GERP scores in any population-trait types.

In contrast to our results in maize, pleiotropy is pervasive in human and mouse literature [28–30,44,54]. It has been shown that highly pleiotropic loci are associated with changes in gene expression in mice [31] and humans [28], broad expression across many tissues [26,29], and an enrichment in accessible chromatin [26]. These enrichment patterns may not be present in maize due to constant selective pressures on novel pleiotropic loci originating after its domestication from teosinte roughly 9,000 years ago [55,56] and constant selection in modern breeding programs, making pervasive wide- and large-effect pleiotropic loci big targets for selection. In a population with large standing natural variation, mutations within wide- and large-effect loci would not be advantageous. This agrees with Fisher and Orr models where mutations causing larger phenotypic effects are less likely to be beneficial over small effect mutations and thus more likely to affect complex organisms [57,58]. Additionally, genetic independence among traits is essential so that favorable combinations of alleles can aid in the adaptation of novel varieties suited to new environments and conditions [59]. Thus, our results of modest pleiotropy agree with the notion that many independent loci control a single trait, and these loci originate from sites with varying chromatin accessibilities, RNA expression values, and conservation levels.

Trait selection heavily influences pleiotropy results. Small subsets of measured traits originating from the same phytomer or highly correlated traits such as amino acid content [60] or grain quality [61] may already share similar regulatory pathways leading to enrichment in pleiotropic loci [23]. This may lead to unrealistic expectations of the pervasiveness of pleiotropy genome-wide. Because pleiotropy estimates rely heavily on allele frequencies, allele effect sizes, the traits under investigation, mapping resolution, and the selection of causal markers, we argue that many claims of pleiotropy in the plant literature are misleading. Additionally, pleiotropic calls on QTL intervals in prior studies in wheat [62], rice [61,63], maize [34,35], and soybean [64] lack the genomic resolution to distinguish between tight linkage and true pleiotropy due to limited markers and large linkage groups.

Although many pleiotropy claims are misleading, this does not mean that pleiotropy does not exist in plants. These pleiotropic relationships may be more confined to fixed, large-effect loci such as *teosinte branched 1* (*tb1*), *vegetative transition 1* (*vgt1*), or closely related traits that share similar developmental pathways. These closely related traits exhibing some pleiotropy include key regulators of flowering time between days to silking, days to anthesis, and the anthesis-silking interval [3], some inflorescence architecture traits [16], and co-regulation between flowering time and leaf number [16,19,20].

In this study, we achieved phenotypic and marker saturation to assess the prevalence of pleiotropy within the maize genome. There is minimal pleiotropy present that, in aggregate, shows little to no relationship with gene expression, sequence conservation, or DNA accessibility and is highly impacted by mapping resolution terms including linkage disequilibrium and genomic interval size. Pleiotropy can still exist within the maize genome but is highly constrained by selection and tight linkage of multiple causal loci. We suggest that in future work in the plant genetics community, caution should be taken before claiming pleiotropy is the causal agent controlling multiple phenotypes. We recommend: 1) that mapping and marker resolution is sufficient to distinguish between tight linkage and pleiotropy, 2) that there be robust testing of a null hypothesis of no pleiotropy through statistical methods or null distributions [36,65], 3) that there be enough unrelated traits to create pleiotropy estimates, and 4) that there be functional characterization or validation of pleiotropic claims through the identification of shared biological pathways or fine-mapping. Additionally, there should be more specificity in the types of pleiotropy under investigation by adhering to previously described nomenclature [2,66].

## Materials and Methods

### Collection of a wide range of phenotypes in NAM and GAP

To determine the degree of pleiotropy in the maize genome, we curated traits measured across two maize mapping populations, the US Nested Association Mapping population (US-NAM, or NAM) [67] and the Goodman Association Panel (GAP) [68]. NAM is a biparental population of 26 diverse inbred lines crossed to a common parent, B73, with 5186 progeny, while the Goodman Association Panel consists of 271 diverse inbred lines. Data were collected from 3,873 leaf mass features [37], 116,381 RNA expression values from seven tissues [38], and 343 field traits. Field traits included flowering time [3,69], plant architecture [21,46,70–75], disease and insect resistance [76–83], and kernel composition [84–90]. Many of these traits are replicated across environments, so a single representative version of that trait was kept. These traits, their publication of origin, and trait values are available in S1-S3 Tables. Due to their size, GAP mass features and expression data are provided in their original publications [37,38].

As shown in the visual methods, we split all phenotypes into four categories based on population and trait type (Fig 1). These population-trait categories were: 1) NAM field, 2) GAP field, 3) GAP mass features, and 4) GAP expression traits. Some traits within GAP field traits included kernel carotenoids and metabolites [84,86,88–90], which could be classified into GAP mass features type because they are also molecular phenotypes. However, only the metabolites measured within a single study were classified as GAP mass features [37].

To create an estimate of statistical pleiotropy to compare to our estimates of biological pleiotropy, we permuted each phenotype ten times for NAM field, GAP field, and GAP mass features. Due to their large number, we permuted GAP expression traits five times. Phenotypes were permuted by fitting a linear model with their respective principal components or PEER (Probabilistic Estimation of Expression Residuals) factors [91] as described in the association mapping section, and we gathered their fitted and residual values. Residual values were independently permuted five or ten times before adding back in the effects of the fitted values. This permutation step ensured that covariances between traits were broken and that the effect of the principal components and PEER factors did not inflate GWA p-values due to their underlying population structure. The genotype matrix was not permuted to maintain the LD structure between SNPs and QTL.

### Association mapping of a wide range of phenotypes

We used the previously generated NAM and GAP Hapmap 3.2.1 SNPs imputed using Beagle 5.1 [92] with the USDA-ARS NCRPIS collection SNPs as the reference [93]. After filtering based on a minimum allele frequency of 0.01, roughly 17 million and 15 million SNPs remained in the GAP and NAM populations, respectively.

GAP field and mass feature traits were mapped with the following model to account for population structure: *y* = *Xβ*_*1*_ + *3 global PCs* + *e*. The three global principal components (PCs) were calculated on a subset of 66,527 SNPs from 3,545 diverse inbred lines in the USDA-ARS NCRPIS collection, and then GAP SNPs were rotated to such PCs. The 66,527 SNPs were chosen because they represented sites with no missing data shared between the three maize panels. This method offered similar control for population structure and kinship over PCs calculated solely within GAP and was consistent when mapping traits across multiple populations [94]. This also saves on computational time compared to using mixed linear models. For GAP expression traits, twenty-five tissue-specific PEER factors were added to the model to control for experimental design and sampling variability as described previously [38].

Due to the population and family structure arising from NAM’s half-sib design, more stringent control of population structure was needed to reduce the false-positive rate. The following two models were used in NAM: *y* = *Xβ*_*1*_ + *3 global PCs* + *main window PCs* + *e* and *y* = *Xβ*_*1*_ + *3 global PCs* + *mid window PCs* + *e*, where the global PCs were calculated similarly to GAP but using NAM SNPs. Main window PCs were calculated only for NAM by breaking the genome into 360 gene windows based on B73 AGPv4 gene coordinates, and PCs were calculated within each window. A total of 360 genes were chosen per window because they offered a reasonable number of degrees of freedom when mapping within NAM. For each window, enough PCs were taken to explain 15% of the total variance. To combat potential edge effects arising from these ‘main’ windows, we slid the windows over by 180 genes, recalculated window PCs (named ‘mid’ window PCs), and then used both collections of PCs in the association analysis. This resulted in 469 ‘main’ window PCs and 512 ‘mid’ PCs. The union of significant sites between these two models was used for further analyses. In cases where both models had a significant association for a single SNP, the most significant association (with the smallest p-value) was taken. All PCs used to map traits in NAM and GAP are available in S4 Table. All observed data (not permuted) and permuted phenotypes were then mapped using the rTASSEL package in R [95] with the Fast Association method [96], keeping all trait-SNP associations with p < 0.00001.

### Calculating a quantitative pleiotropy score

Due to high linkage disequilibrium and the difficulty in ascertaining the true causal SNP(s) from a list of significant associations, we chose to investigate pleiotropy within genic and intergenic intervals. We did this by breaking the genome into an equal number of genic and intergenic intervals based on gene annotations in B73 AGPv4 with the GenomicRanges package in R [97]. Any overlapping genic ranges were merged into one large interval. This resulted in n = 75,490 intervals with a mean length of 27,902 bp.

Using a custom TASSEL plugin (version 5.2.79, CountAssociationsPlugin) [98], we counted the number of trait-associated SNPs in each interval for each population-trait type separately for both the observed and permuted results. The matrices of intervals by trait counts were then used to calculate the number of unique (non-redundant) traits mapping to each interval. We chose to calculate the number of traits mapping to each interval over using other previously established pleiotropy methods because it provided a quantitative score on the variability of trait-SNP associations, made no assumptions on which single SNP was the ‘most’ pleiotropic, did not require effect sizes of SNPs, and provided an estimate of how pleiotropic a specified interval was rather than a single SNP.

Investigation of pleiotropy enrichment compared to the permuted data was performed by taking the log of the observed divided by the mean permuted pleiotropy count for each interval plus of peseudo-count of plus one. The pseudo-count of one in the numerator and denominator ensured that all intervals were retained in the plot.

### Functionally characterizing pleiotropy using linear and random forest models

To determine what factors may impact pleiotropy in maize, we investigated which annotations were the most important in determining the degree of pleiotropy within our traits and populations. We trained three random forest models using the R ranger package [99] and one gradient boosting model using the R XGBoost package [100], using features that are known to statistically interfere with GWA analyses and features that may biologically impact pleiotropy.

Model 1 was a random forest model that only included nuisance (mapping) features. All terms were calculated for each interval and included average R^2^ linkage disequilibrium within NAM and GAP calculated with the MeanR2FromLDPlugin function, the total number of SNPs per interval used for the GWA analysis before p-value filtering, and the interval size. All features were calculated for each population separately.

Models 2 (random forest) and 4 (gradient boosting) included both nuisance and biological features. Biological features included the mean Genomic Evolutionary Rate Profiling (GERP) scores of GERP SNPs that overlapped exactly (based on their physical position) with significant GWA SNPs averaged across an interval, the number of ATAC-seq sites previously analyzed [42], RNA and protein expression values in B73 across 23 diverse tissues [40], the interval type (i.e., genic or intergenic), and adjusted GAP expression pleiotropy values (described below).

For Model 3, we included all nuisance and biological variables as Models 2, and 4; however, we also added fourteen biological process or molecular function gene ontology terms to the model. These terms describe the presence or absence of that ontology term to that interval. These gene ontology terms were scored for each interval where 1 denoted that genic interval contained a gene belonging to that gene ontology term, while 0 meant that genic and all intergenic intervals did not harbor a gene belonging to that ontology term. These terms included transcription DNA-templated (BP, GO:0006351), translation (BP, GO:0006412), regulation of transcription DNA-templated (BP, GO:0006355), regulation of translation (BP, GO:0006417), leaf development (BP, GO:0048366), regulation of leaf development (BP, GO:2000024), flower development (BP, GO:0009908), regulation of flower development (BP, GO:0009909), regulation of catalytic activity (BP, GO:0050790), signal transduction (BP, GO:0007165), nucleotide binding (MF, GO:0000166), DNA binding (MF, GO:0003677), RNA binding (MF, GO:0003723), and DNA-binding transcription factor activity (MF, GO:0003700).

All models were run individually for each population-trait type (eight separate versions of each model, 32 total models), training the model on data from chromosomes 1–9 and holding out chromosome 10 for model testing and validation. The random forest models (Models 1-3) trained using R/ranger were built with default parameters, using the impurity variable importance mode and 500 trees. For the gradient tree boosting model (Model 4) we performed a grid search over the hyperparameter space of eta and max.depth and chose the values eta = 0.5, max.depth = 6 for each population, as these values attained approximately the lowest test error in each population. All other XGBoost parameters were set to their default values. Each model’s relative importance scores were calculated by dividing all importance scores by the maximum importance score. A correlation matrix of all model terms is available in S1 Fig.

Due to the high importance of linkage disequilibrium, the number of input SNPs, and interval size in the random forest models, an adjusted pleiotropy metric was created. Adjusted pleiotropy was calculated by subtracting the predicted values of the following random forest model: *pleiotropy ∼ averageLinkageDiseqilibrium + numberInputSNPs + intervalSize* (Model 1) from the observed pleiotropy for each population-trait type. The relationship between adjusted pleiotropy and observed pleiotropy is available in S8 Fig. Associations with adjusted pleiotropy were then assessed for the max B73 expression across numerous tissues and the average GERP scores using linear models in R. These adjusted pleiotropy values aimed to control for spurious pleiotropy, however, we cannot distinguish between instances of true biological, mediated, or spurious pleiotropy. All raw and adjusted pleiotropy values and biological annotations used for analyses are available in Supplemental Table 6.

### Enrichment of adjusted pleiotropy scores with gene ontology terms

To investigate which gene ontology terms were enriched in highly and lowly pleiotropic regions, we subsetted our adjusted pleiotropy scores to genic regions, uplifted our B73 gene models from AGPv4 to RefGen_v5, and connected the RefGen_v5 genes with GO terms using Zm-B73-REFERENCE-NAM-5.0_Zm00001eb.1.interproscan.tsv obtained from MaizeGDB [101]. We then gathered the top twenty-five percent and bottom twenty-five percent of genic above zero adjusted pleiotropy values in each population-trait type and labeled these loci as highly pleiotropic and lowly pleiotropic, respectively. We then used the intervals in these percentiles as inputs into topGO [102]. We investigated the genes associated with molecular function, biological processes, and cellular components, retaining all significant false discovery rate (FDR) corrected terms from a Fisher’s Exact test (p < 0.05). Significant GO terms and all other plots in this manuscript were visualized using ggplot2 [103].

## Code and Data Availability

All phenotypic data for this project can be found in the supplementary information tables and all code for this project can be found on Bitbucket (https://bitbucket.org/bucklerlab/pleiotropy/src/master/).

## Acknowledgments

We thank Sara Miller for copy editing and Michelle Stitzer for their suggestions in the preparation of this manuscript. This work was supported by the USDA NIFA AFRI predoctoral fellowship (MBKB: 2022-67011-36458; TF: 2022-67011-36564), the USDA-ARS, and the NSF PGRP (IOS#1822330).

## Supplemental information captions

S1 Tab1e: Publications and traits used in this study.

S2 Table: NAM phenotypic field data used in the analysis.

S3 Table: GAP field data used in the analysis.

S4Table: Global, main window, and midway window principal components used to map NAM traits and global principal components used to map phenotypes in GAP.

S5 Table: Random forest and gradient boosting model performances and prediction accuracy for the four population-trait categories for the observed and permuted data. R^2^ prediction accuracy is calculated on held-out chromosome 10 interval data from a model trained using data from chromosomes 1 through 9. Prediction accuracy for models 1 (random forest, only nuisance terms), 2 (random forest), 3 (random forest with gene ontology terms), and 3 (gradient boosting) are provided.

S6 Table: Aggregated raw pleiotropy counts, adjusted pleiotropy counts, and values for all biological features used for analyses.

S7 Table: Top twenty intervals and their associated genes by population and trait type.

## Supplemental Figures

**S1 Fig:**
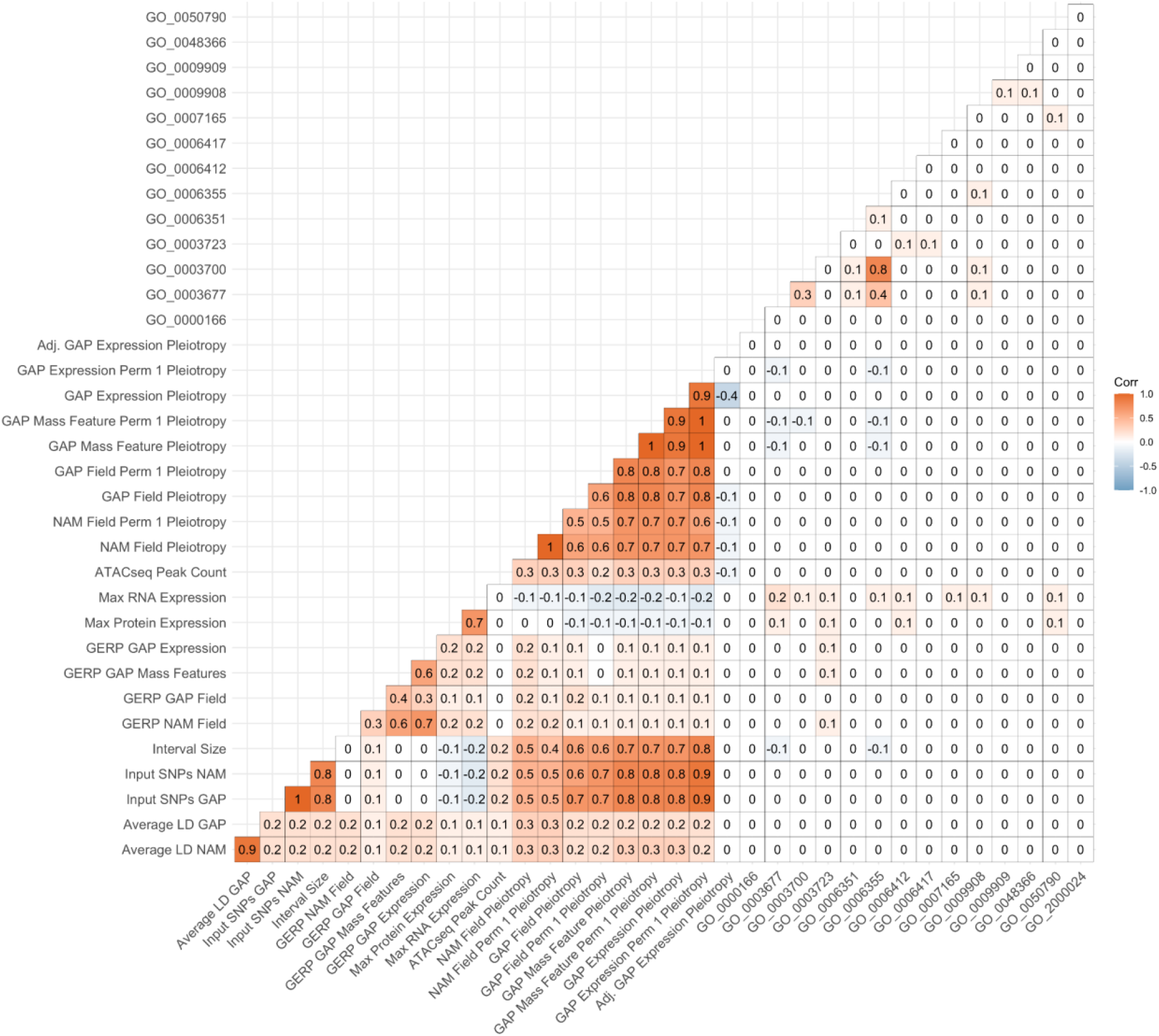
Correlations between terms used within the random forest and gradient boosting models. Only the first permuted value from each of the four-population trait categories was included in the correlation matrix for simplicity.

**S2 Fig:**
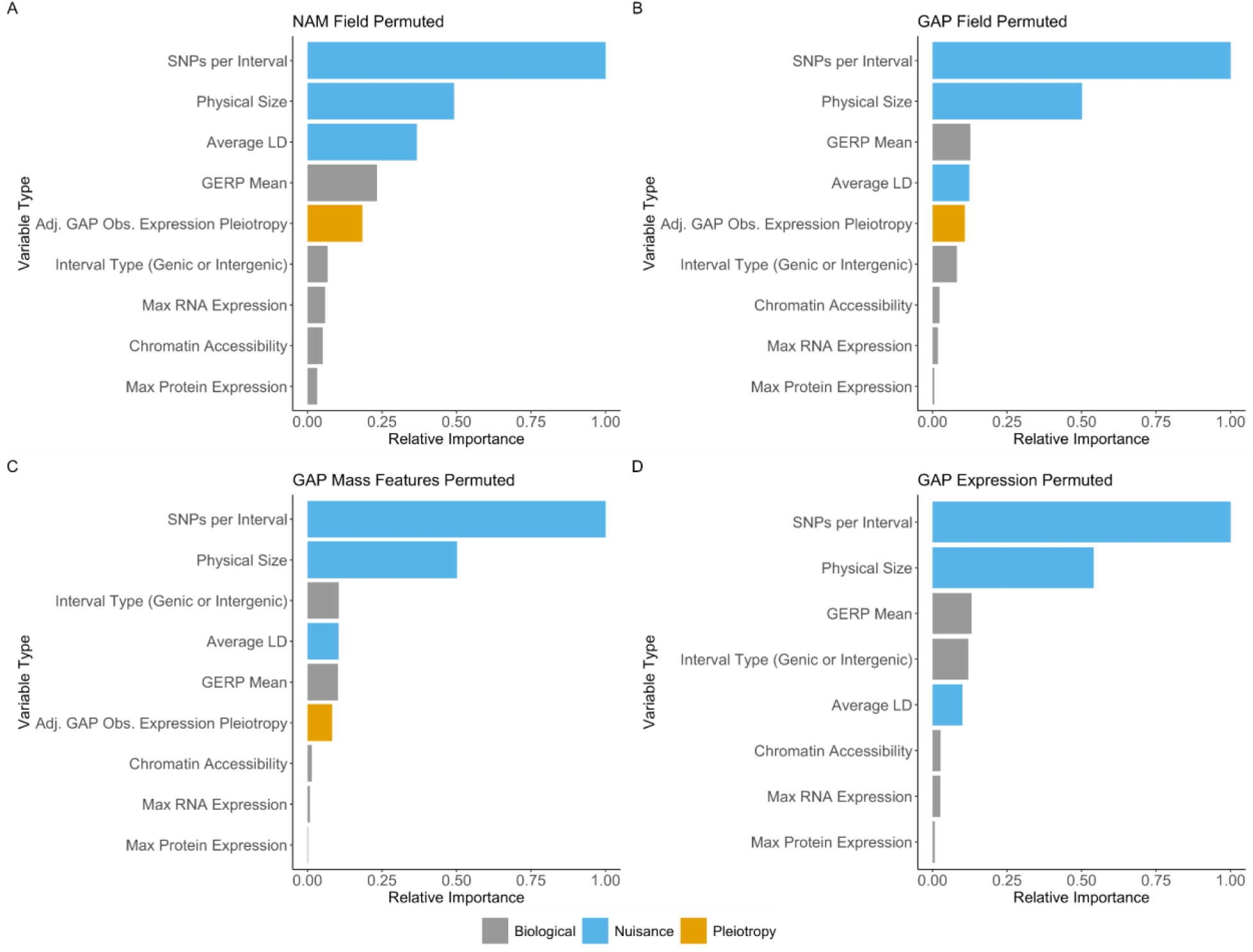
Relative importance scores from random forest models of biological and nuisance features in explaining the permuted pleiotropy values for Model 2 (random forest without gene ontology terms). Across all four population-trait categories, nuisance variables results showed higher relative importance over biological features. The plots show the observed (not permuted) data for the (**a**) NAM field, (**b**) GAP field, (**c**) GAP mass features, and (**d**) GAP expression data.

**S3 Fig:**
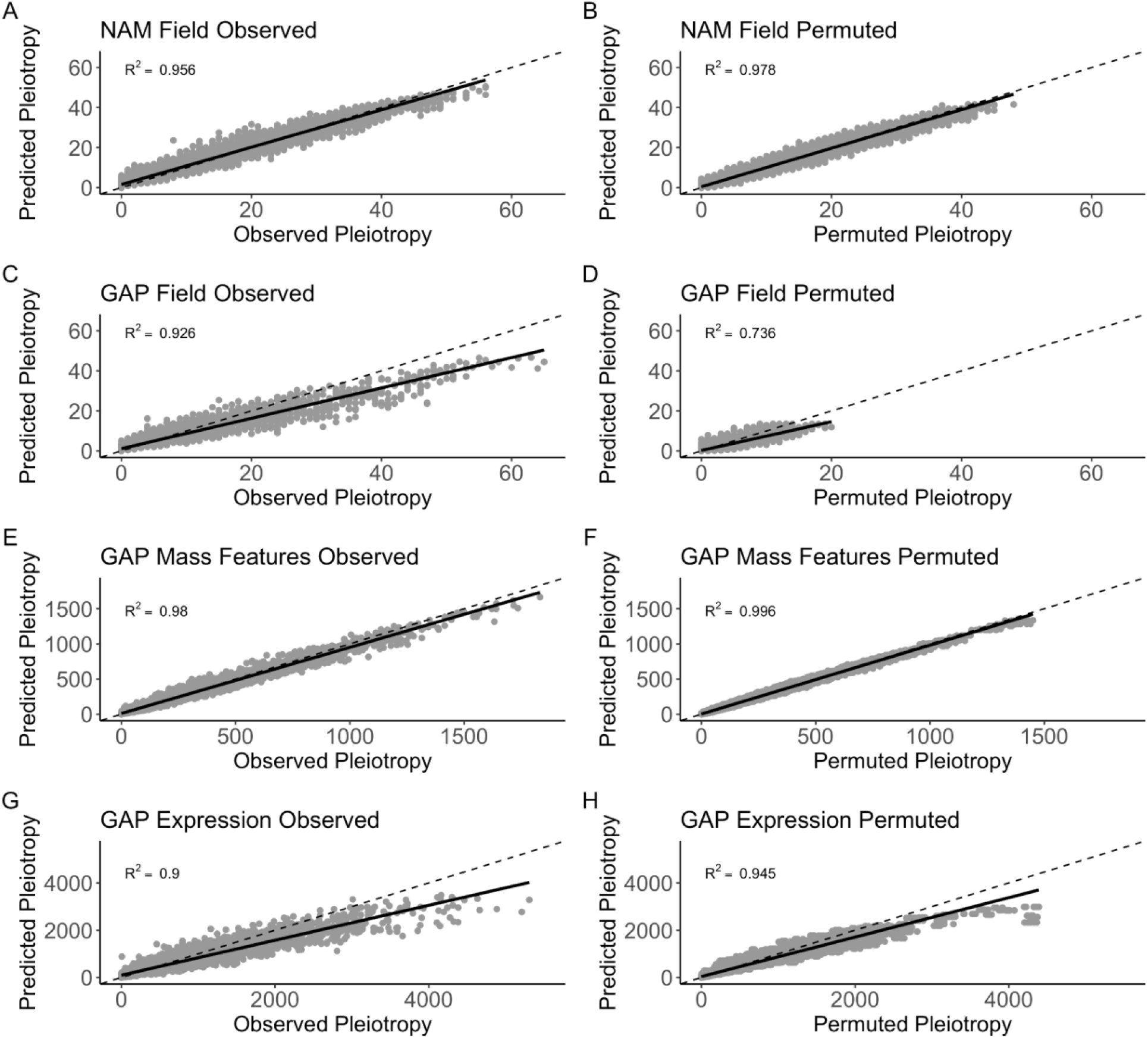
Predicted versus observed pleiotropy values for Model 2 (random forest without gene ontology terms) from eight individual random forest models trained on pleiotropy values from chromosomes 1-9 and tested on chromosome 10 data. The dashed line represents the 1-1 identity line, while the solid line represents fitted values. Panels (a), (c), (e), and (g) show the observed (not permuted) results while panels (b), (d), (f), and (h) show the permuted results. Panels (a) and (b) show NAM field, (c) and (d) GAP field, (e) and (f) GAP mass features, and (g) and (h) GAP expression.

**S4 Fig:**
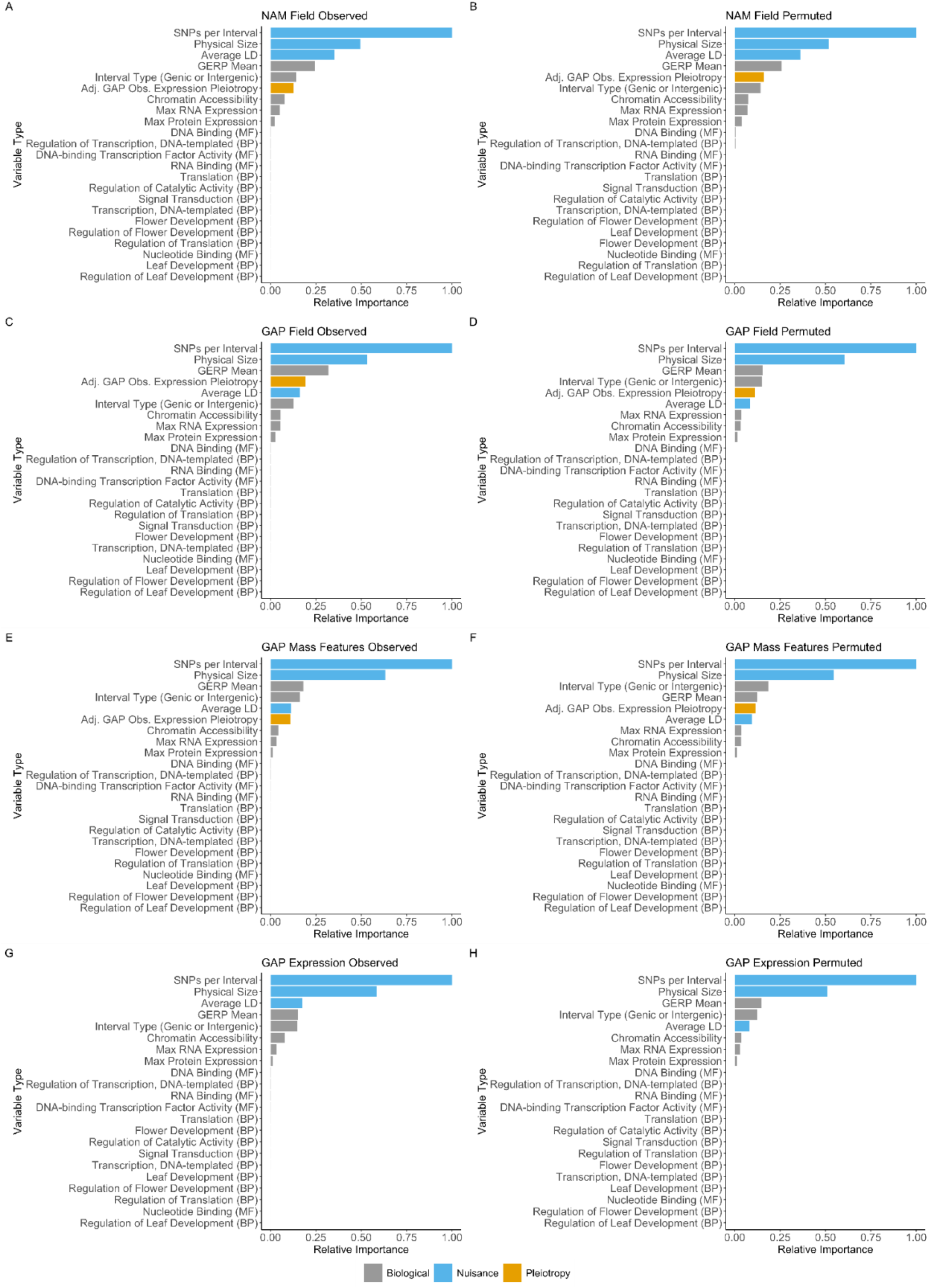
Relative importance scores from random forest models of biological, nuisance, and gene ontology features in explaining the observed and permuted pleiotropy values for Model 3 (random forest with gene ontology terms). Across all four population-trait categories, nuisance variables results showed higher relative importance over biological features. The plots show data for the observed data in panels (a), (c), (e), and (g) and the permuted data in panels (b), (d), (f), and (h). Panels (a) and (b) show NAM field results, (c) and (d) GAP field, (e) and (f) GAP mass features, and (g) and (h) GAP expression data.

**S5 Fig:**
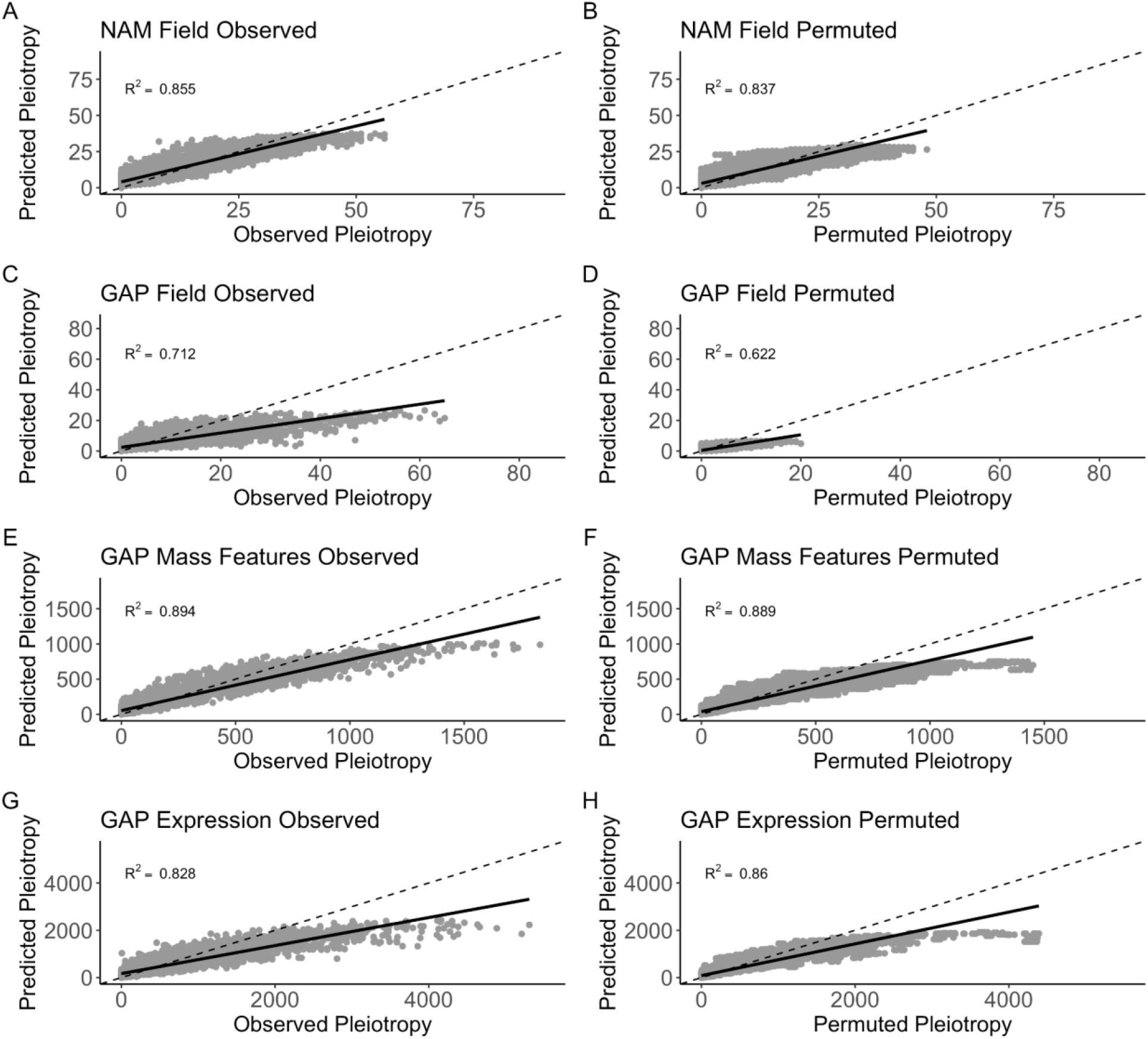
Predicted versus observed pleiotropy values for Model 3 (random forest with gene ontology terms)from eight individual random forest models trained on pleiotropy values from chromosomes 1-9 and tested on chromosome 10 data. The dashed line represents the 1-1 identity line, while the solid line represents fitted values. Panels (a), (c), (e), and (g) show the observed (not permuted) results while panels (b), (d), (f), and (h) show the permuted results. Panels (a) and (b) show NAM field, (c) and (d) GAP field, (e) and (f) GAP mass features, and (g) and (h) GAP expression.

**S6 Fig:**
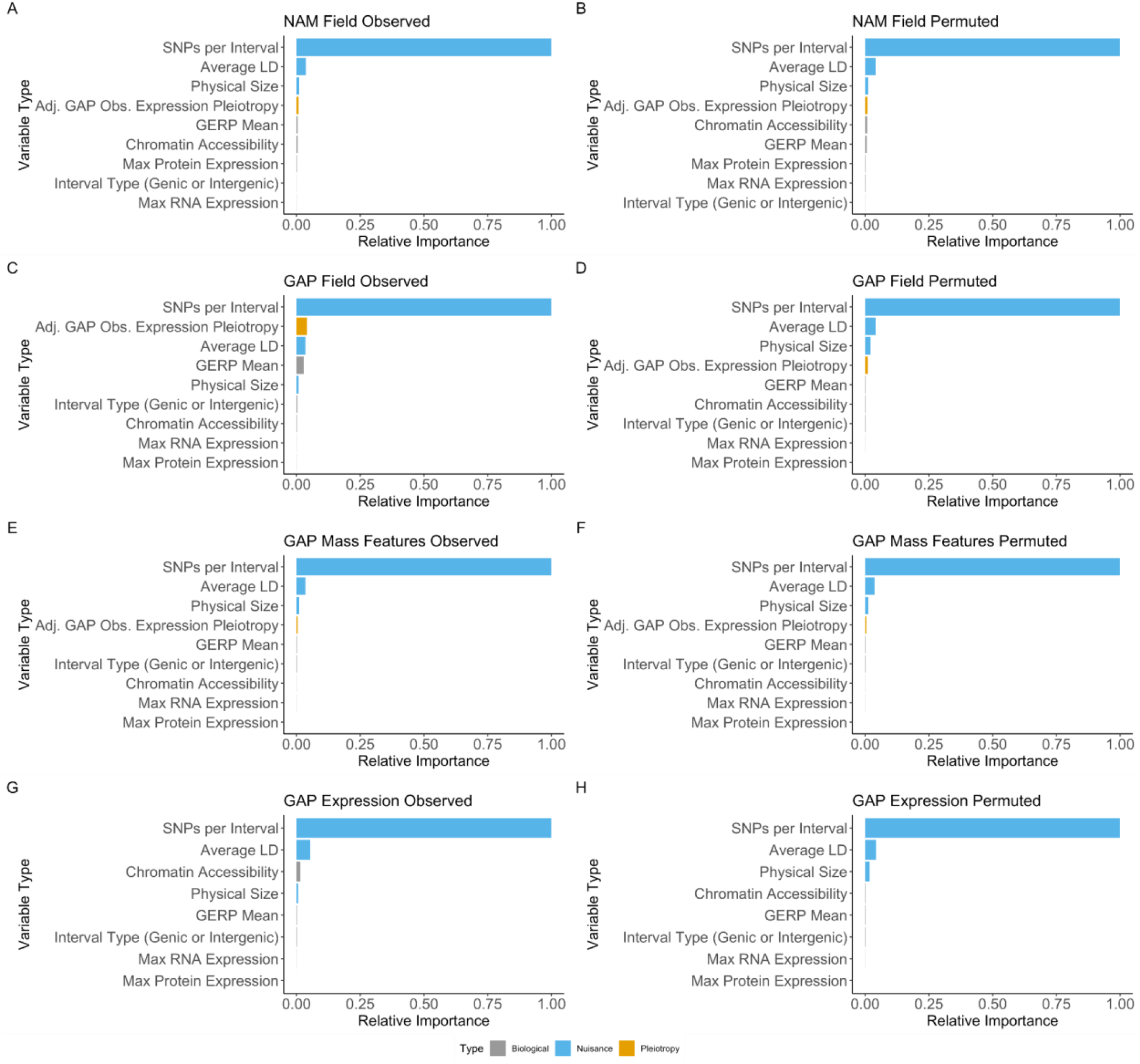
Relative importance scores from random forest models of biological, nuisance, and gene ontology features in explaining the observed and permuted pleiotropy values for Model 4 (gradient boosting). Across all four population-trait categories, nuisance variables results showed higher relative importance over biological features. The plots show data for the observed data in panels (a), (c), (e), and (g) and the permuted data in panels (b), (d), (f), and (h). Panels (a) and (b) show NAM field results, (c) and (d) GAP field, (e) and (f) GAP mass features, and (g) and (h) GAP expression data.

**S7 Fig:**
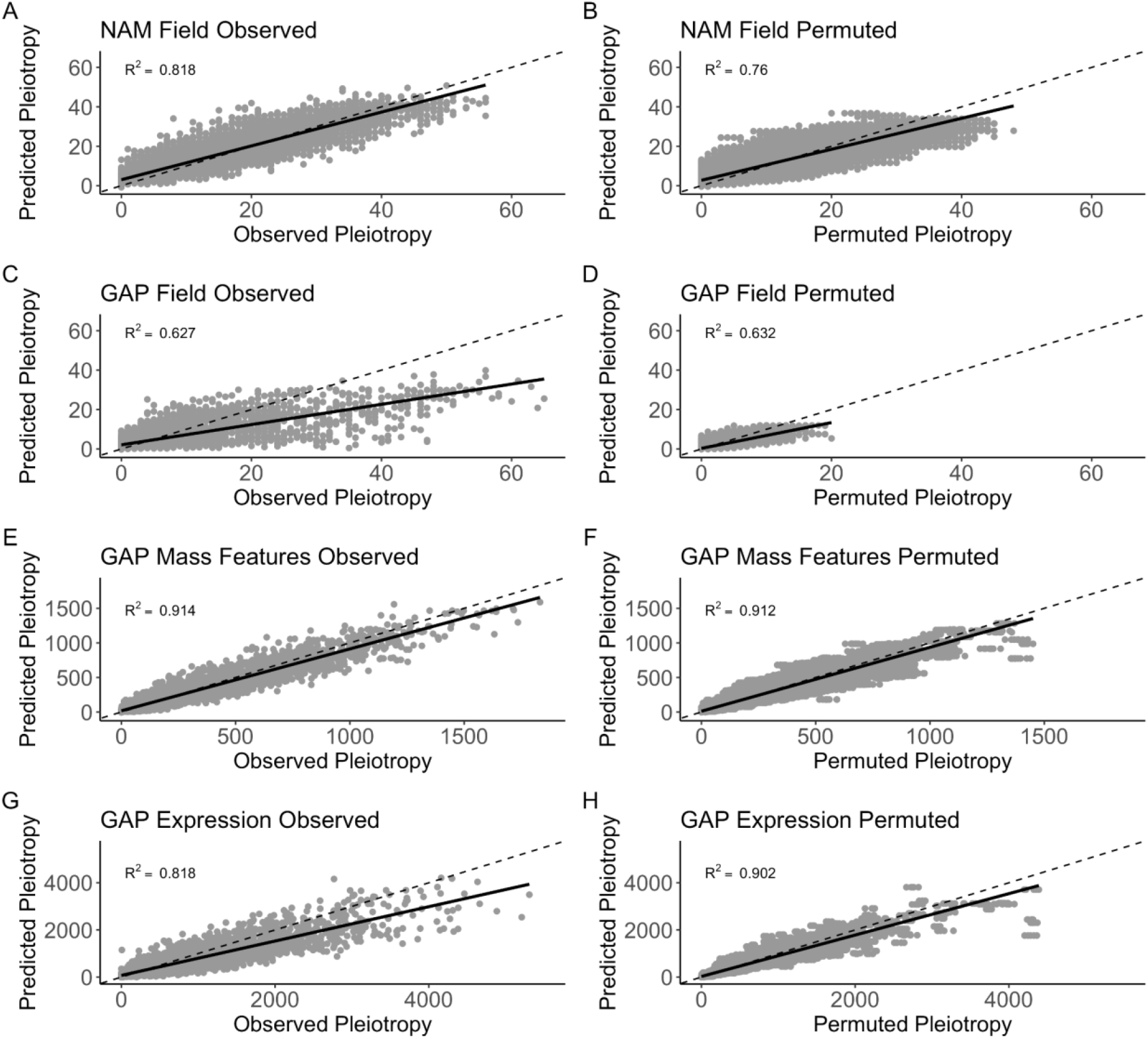
Predicted versus observed pleiotropy values for Model 4 (gradient boosting) from eight individual random forest models trained on pleiotropy values from chromosomes 1-9 and tested on chromosome 10 data. The dashed line represents the 1-1 identity line, while the solid line represents fitted values. Panels (a), (c), (e), and (g) show the observed (not permuted) results while panels (b), (d), (f), and (h) show the permuted results. Panels (a) and (b) show NAM field, (c) and (d) GAP field, (e) and (f) GAP mass features, and (g) and (h) GAP expression.

**S8 Fig:**
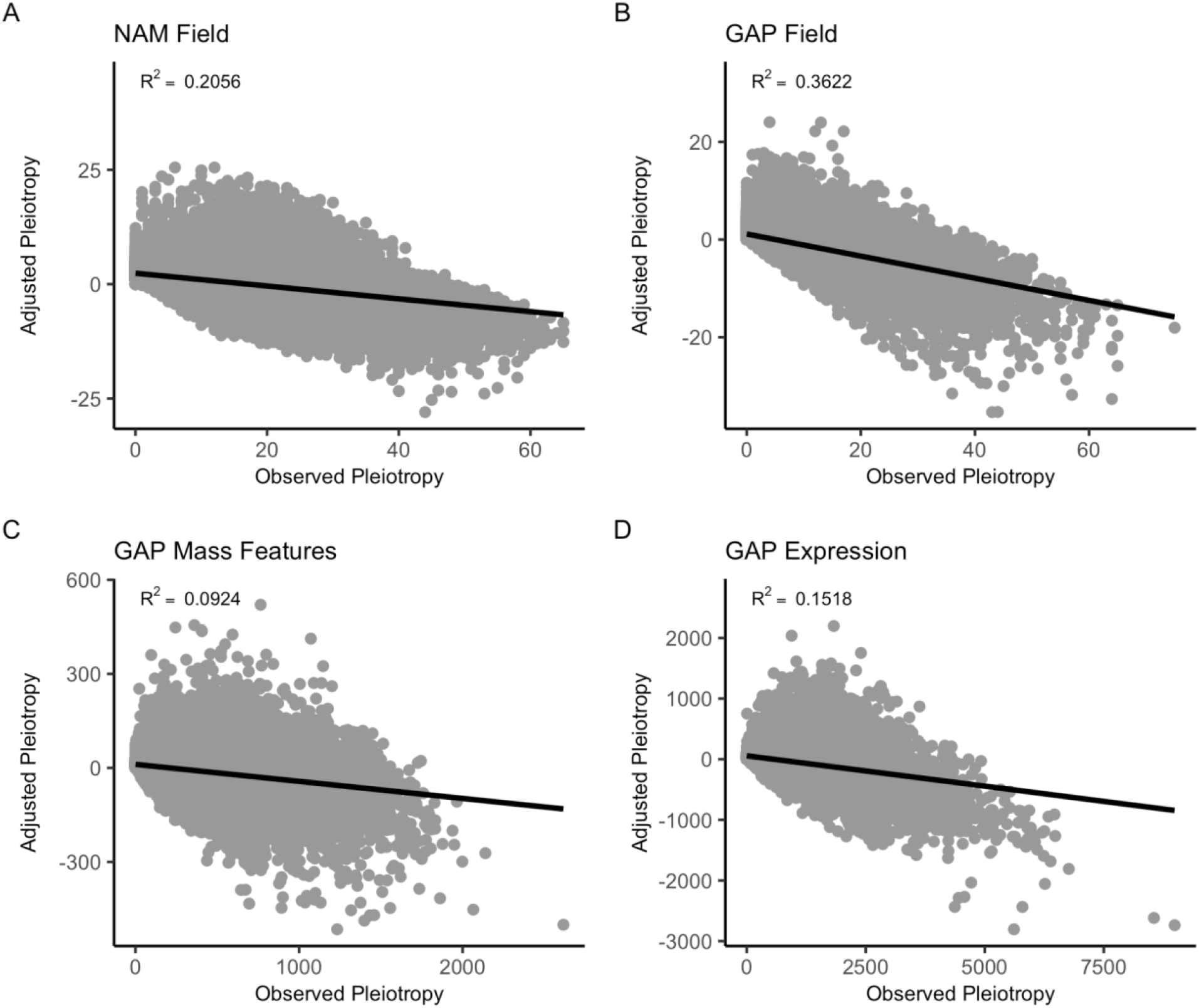
There is a small, negative relationship between adjusted pleiotropy and observed pleiotropy in the four population-trait types. The scatter plots show the relationship between the adjusted and observed pleiotropy values for (a) NAM field, (b) GAP field, (c) GAP mass features, and (d) GAP expression traits. Values in the top right of each plot show the R^2^ values from correlating the adjusted versus unadjusted pleiotropy data.

**S9 Fig:**
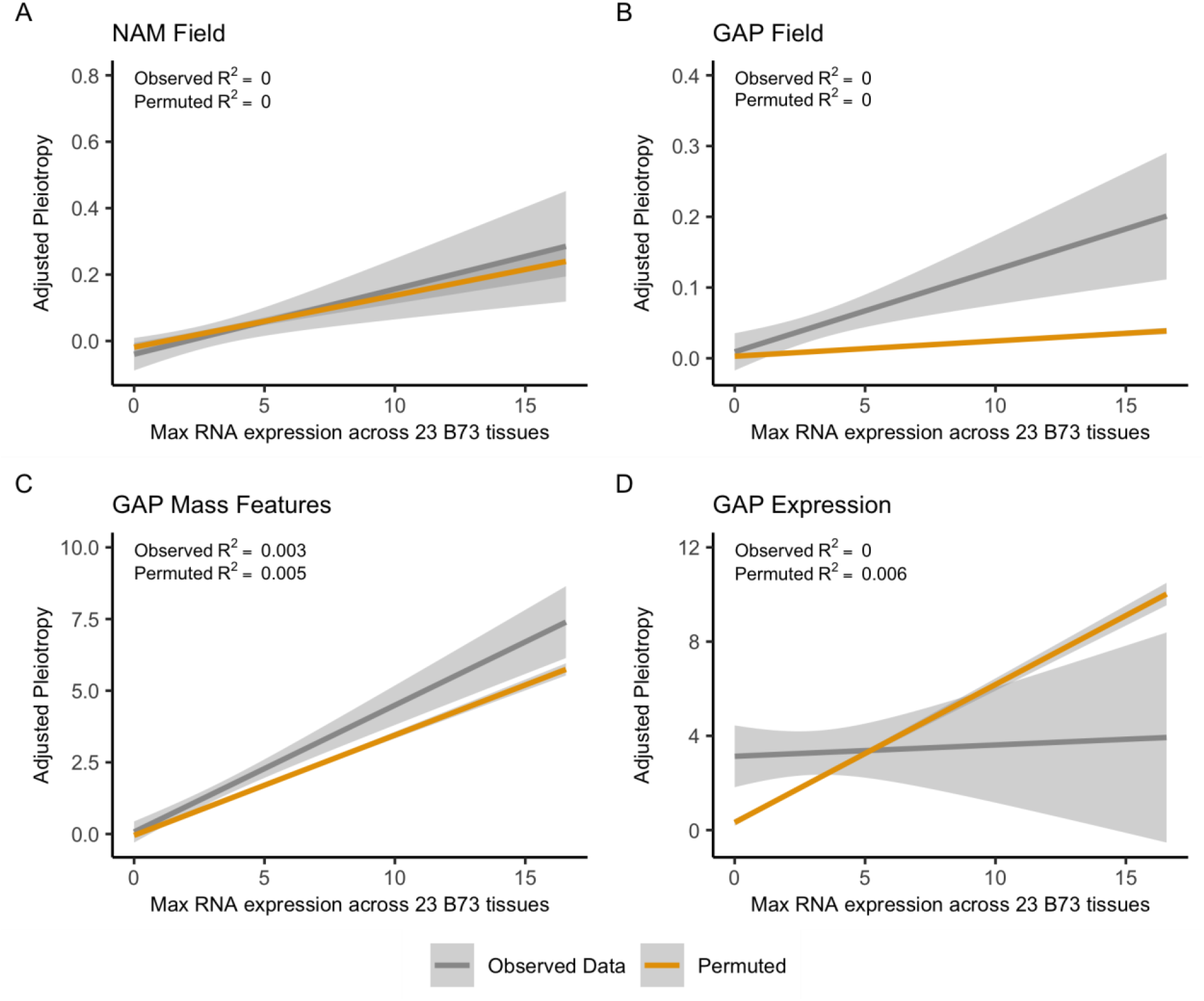
Adjusted pleiotropy is not strongly correlated with the max B73 RNA expression across 23 diverse tissues. The line plots of adjusted pleiotropy for only genic ranges against the max RNA expression show the observed (not permuted, gray lines) and permuted (yellow lines). Panels show adjusted pleiotropy scores for (a) NAM field, (b) GAP field, (c) GAP mass features, and (d) GAP expression traits. Values in the top right of each plot show the R^2^ values from correlating the observed genic and permuted genic adjusted pleiotropy data separately against the max RNA expression value across 23 B73 tissues. Shading around lines shows the 95% confidence interval.

**S10 Fig:**
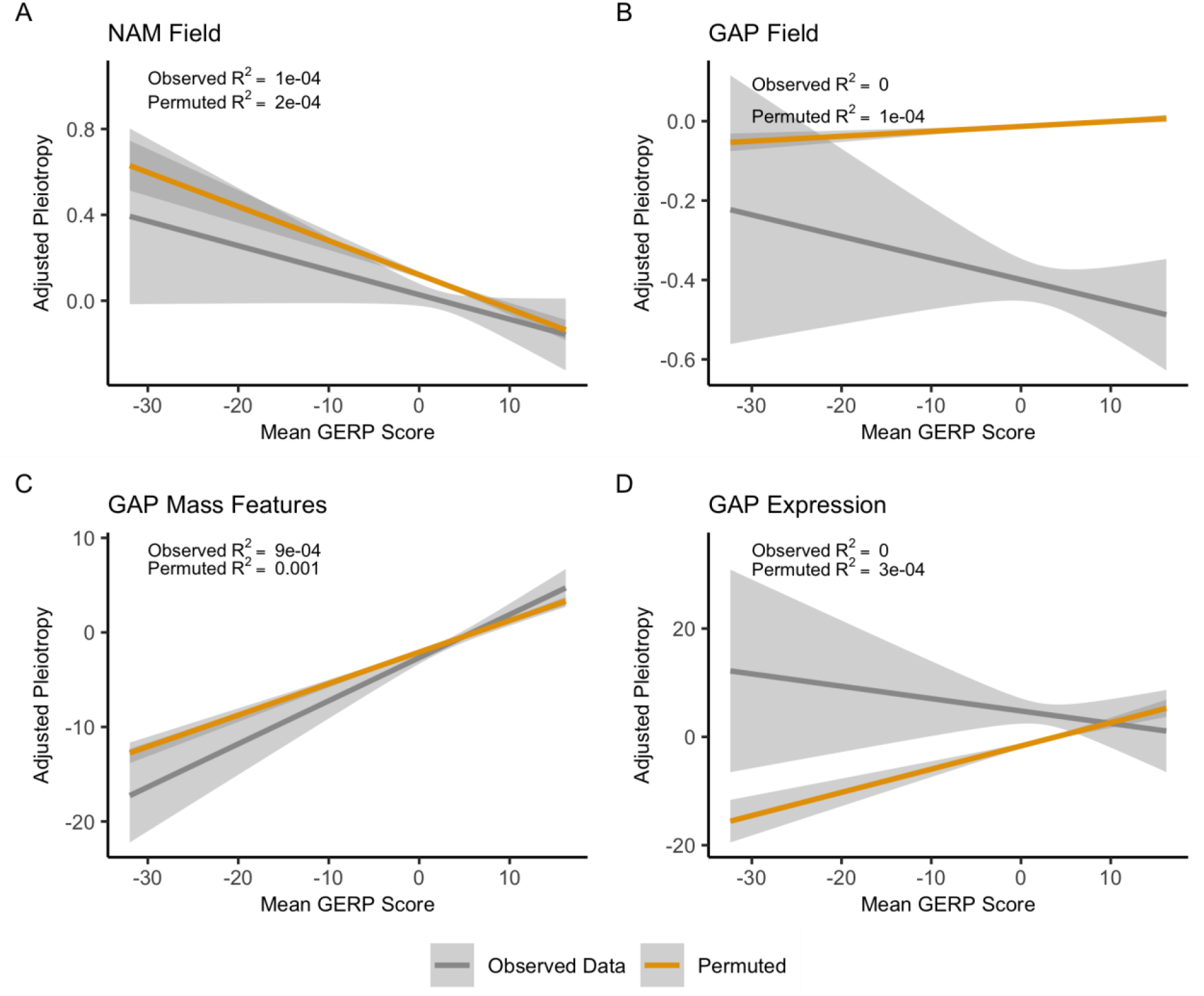
Adjusted pleiotropy is not correlated with the mean GERP score. Line plots showing the adjusted pleiotropy against the mean GERP score for all genic and intergenic intervals. Mean GERP was calculated separately by averaging overlapping observed (not permuted) GWA SNPs with GERP SNPs for each population-trait type. Results for the observed data are in gray, while permuted results are in yellow. Panels show adjusted pleiotropy scores for (a) NAM field, (b) GAP field, (c) GAP mass features, and (d) GAP expression traits. Values in the top right of each plot show the R^2^ values from correlating the observed and permuted adjusted pleiotropy data separately against the mean GERP score. Shading around lines shows the 95% confidence interval.

